# Lyophilized Extracellular Vesicles Retain Regenerative Activity and Accelerate Wound Healing

**DOI:** 10.64898/2026.06.10.731475

**Authors:** Young-Ju Lim, Axel Schmitter-Sánchez, Min-Soo Seo, Gun Woo Lee, Min-Jung Ma, Wansuk Son, Seunghyun Kang, Joo-Hee Choi, Wook Tae Park, Sangbum Park

## Abstract

Extracellular vesicles (EVs) are promising therapeutic agents for tissue regeneration because they regulate intercellular communication and modulate inflammatory responses. However, preserving EV bioactivity during long-term storage remains a major barrier to clinical application. We examined whether lyophilized EVs stored at −80 °C maintain their structural integrity and therapeutic efficacy in an in vivo wound-healing model. Mesenchymal stem cell-derived EVs were isolated and freeze-dried before storage at −80 °C. We evaluated EV physicochemical characteristics before and after lyophilization using nanoparticle tracking analysis, transmission electron microscopy, and EV marker-expression analysis. To assess regenerative efficacy, lyophilized EVs were applied topically to full-thickness ear wounds in CCR2-GFP mice. Wound-healing progression and CCR2-positive cell infiltration were monitored during tissue recovery using intravital microscopy. Lyophilized EVs preserved their characteristic morphology, particle-size distribution, and EV surface marker expression after storage. In vivo analysis showed that EV-treated wounds closed significantly faster than phosphate-buffered saline-treated controls. Additionally, lyophilized EV treatment reduced CCR2-positive inflammatory cell recruitment during healing, suggesting an immunomodulatory role in tissue regeneration. These findings show that EVs lyophilized and stored at −80 °C retain biological function and therapeutic potential in vivo. Lyophilized EVs may, therefore, provide a practical strategy for long-term storage and delivery of EV-based regenerative therapeutics.

## 1. Introduction

Extracellular vesicles (EVs) are nanosized membrane-bound particles secreted by various cell types that play critical roles in intercellular communication through the transfer of proteins, lipids, and nucleic acids (1). In recent years, EVs have gained considerable attention as promising therapeutic agents in regenerative medicine owing to their ability to modulate inflammation, promote tissue repair, and mediate paracrine signaling without the limitations associated with direct stem cell transplantation (2). Among the various sources of EVs, mesenchymal stem cell (MSC)-derived EVs have shown substantial therapeutic potential in a wide range of tissue injury and wound healing models (3, 4).

Despite their therapeutic promise, several limitations remain regarding the clinical application of EV-based therapies. One of the major challenges is maintaining EV stability and biological activity during long-term storage. Conventional preservation methods, including repeated freeze–thaw cycles, may induce vesicle aggregation, membrane disruption, and functional degradation, thereby reducing therapeutic efficacy (5-8). Therefore, establishing an efficient preservation strategy capable of maintaining EV integrity and functionality is essential for the clinical translation and commercialization of EV therapeutics.

Lyophilization, also referred to as freeze-drying, has emerged as a promising approach for improving the long-term stability and storage efficiency of biological materials (9). By removing water under low-temperature and vacuum conditions, lyophilization minimizes structural damage and may prolong the shelf life of EV preparations (7). Previous studies have suggested that lyophilized EVs can preserve their physicochemical characteristics under appropriate storage conditions (7). However, evidence demonstrating whether freeze-dried EVs retain their in vivo regenerative efficacy after storage remains limited, particularly in tissue repair and wound healing models (5).

Wound healing is a complex biological process involving coordinated inflammatory responses, immune cell recruitment, angiogenesis, and extracellular matrix remodeling (10). Among inflammatory cell populations, CCR2-positive monocytes and macrophages are known to play critical roles during the inflammatory and regenerative phases of tissue repair. Dysregulated or prolonged inflammatory responses can impair wound healing, whereas controlled modulation of inflammatory cell recruitment may enhance tissue regeneration (11). Therefore, understanding the effects of EV treatment on CCR2-positive inflammatory cell dynamics may provide important insights into the immunomodulatory mechanisms underlying EV-mediated tissue repair (4, 12, 13).

In the present study, we investigated whether lyophilized MSC-derived EVs stored at −80 °C retain their structural integrity and therapeutic efficacy in vivo. EVs were isolated from MSCs, subjected to freeze-drying, and characterized using nanoparticle tracking analysis (NTA), transmission electron microscopy (TEM), and EV surface marker analysis (14). To evaluate their regenerative potential, lyophilized EVs were topically administered to full-thickness wounds on the mouse ear (15). Additionally, wound closure and CCR2-positive inflammatory cell infiltration were visualized in vivo during the healing process. Our findings show that lyophilized EVs preserved under −80 °C conditions maintain their biological functionality and promote tissue regeneration, supporting their potential application as clinically feasible EV-based therapeutics.

## 2. Materials and Methods

### 2.1 Reagents

Dulbecco’s Modified Eagle Medium (DMEM), phosphate-buffered saline (PBS), penicillin-streptomycin (P/S), trypsin-ethylenediaminetetraacetic acid (trypsin-EDTA), fetal bovine serum (FBS), and exosome-depleted FBS were purchased from Gibco-BRL (Carlsbad, CA, USA). StemPro Osteogenesis, Adipogenesis, and Chondrogenesis Differentiation Kits were obtained from Gibco, Thermo Fisher Scientific (Waltham, MA, USA). Alizarin Red S, Oil Red O, Alcian Blue 8GX, formaldehyde, sucrose, and HEPES were purchased from Sigma-Aldrich (St. Louis, MO, USA). Aldehyde/sulfate latex beads were obtained from Thermo Fisher Scientific (Rockford, IL, USA). Formvar carbon-coated copper grids for TEM were purchased from Ted Pella Inc. (Redding, CA, USA).

### 2.2 Cell culture

This study was approved by the Institutional Review Board of Yeungnam University Medical Center (IRB No. 2017-07-032). Human epidural fat tissue was obtained under sterile conditions and minced into small fragments. The tissue was digested enzymatically with collagenase type II (Worthington Biochemical Corporation, Lakewood, NJ, USA) at 37°C for 1 h. After digestion, the cell suspension was filtered through a 70-μm cell strainer and centrifuged to collect the stromal vascular fraction. The isolated cells were cultured as human epidural fat-derived MSCs (hEF-MSCs) in DMEM (Gibco, Carlsbad, CA, USA) supplemented with 10% FBS and 1% P/S. Cells were maintained at 37°C in a humidified atmosphere containing 5% CO2. The culture medium was replaced every other day, and cells were subcultured when they reached approximately 80-90% confluence.

### 2.3 Isolation of EVs

EVs were isolated from hEF-MSC-conditioned medium. To minimize contamination from bovine-derived vesicles, cells were cultured under standard conditions (37°C, 5% CO_2_) in medium supplemented with 10% exosome-depleted FBS (Gibco). Conditioned medium was collected every 48 h, and cellular debris was removed by centrifugation at 300 × g for 10 min. The clarified supernatant was then concentrated using a tangential flow filtration (TFF) system (Pall Corporation, Port Washington, NY, USA) operated at a flow rate of 120 rpm.

### 2.4 Characterization of EVs

MSC surface marker expression was analyzed using flow cytometry (Gallios, Beckman Coulter, Brea, CA, USA) with antibodies against CD90-PE (BioLegend, 555596), CD105-PE (Bio-Rad, Hercules, CA, USA, MCA1557), and CD45-FITC (BioLegend, 555482). EV surface marker expression was evaluated by bead-based flow cytometry (Gallios) using antibodies against CD9 (BioLegend, 358259), CD63 (BioLegend, 353003), and CD81 (BioLegend, 349505). For this analysis, EVs were incubated with 4% aldehyde/sulfate latex beads (Thermo Fisher Scientific, Rockford, IL, USA) for 24 h at room temperature. EV morphology was examined by TEM. EV samples were adsorbed onto formvar carbon-coated copper grids (Ted Pella Inc., Redding, CA, USA), fixed with 2.5% glutaraldehyde for 1 min, air-dried, and observed under a transmission electron microscope. Particle-size distribution and concentration were measured using NTA (NanoSight NS300, Malvern Panalytical, Worcestershire, UK) according to the manufacturer’s instructions.

### 2.5 MSC differentiation assays

hEF-MSCs were seeded in 24-well culture plates for differentiation assays. For osteogenic differentiation, MSCs were cultured with the StemPro Osteogenesis Differentiation Kit (Gibco, Thermo Fisher Scientific, Waltham, MA, USA) for 20 days. After differentiation, cells were fixed with 4% formaldehyde (Sigma-Aldrich, St. Louis, MO, USA) for 5 min at room temperature and stained with 2% Alizarin Red S solution (ARS; Chemicon, Merck Millipore, Billerica, MA, USA) for 15 min. For adipogenic differentiation, MSCs were cultured with the StemPro Adipogenesis Differentiation Kit (Gibco, Thermo Fisher Scientific) for 21 days. Differentiated cells were fixed with 4% formaldehyde for 10 min at room temperature and stained with Oil Red O solution (Sigma-Aldrich) for 15 min. For chondrogenic differentiation, MSCs were cultured in chondrogenic differentiation medium using the StemPro Chondrogenesis Differentiation Kit (Gibco, Thermo Fisher Scientific) for 21 days. After differentiation, the cells were fixed with 4% formaldehyde for 5 min at room temperature and stained with Alcian Blue 8GX (Sigma-Aldrich) for 30 min to assess glycosaminoglycan deposition. MSCs cultured without differentiation supplements served as controls. All differentiation experiments were performed in duplicate and independently repeated three times.

### 2.6 Lyophilization

EVs were collected and concentrated from conditioned medium as described in the preceding sections. Prior to lyophilization, EVs were suspended in a protective solution containing 8.5% (w/w) sucrose and 20 mM HEPES to preserve vesicle stability and biological activity during the freeze-drying process (7, 9, 16). Lyophilization was performed according to a modified protocol based on a previously reported method (7). Briefly, EV suspensions containing the lyoprotectant solution were rapidly frozen, followed by water removal through sublimation under vacuum conditions to maintain vesicle structure and integrity. After lyophilization, EV samples were stored at −80 °C until further use. Before downstream experiments, lyophilized EVs were reconstituted in an appropriate buffer, and their physicochemical properties were evaluated using NTA and EV surface marker analysis.

### 2.7 In vivo wound-healing assay

Mice were anesthetized with vaporized isoflurane, and a full-thickness circular wound was generated on the dorsal ear skin using a 1-mm punch biopsy tool (Miltex, York, PA, USA). Lyophilized EVs were reconstituted in sterile PBS and topically administered to the wound site immediately after injury. Briefly, 4 μL of EV suspension was applied directly to the wound area. Control mice received an equal volume of sterile PBS. For longitudinal wound-healing analysis, the same wound area in each mouse was imaged repeatedly every 3 days after wound induction. Macroscopic wound images were acquired using a Zeiss Stemi 305 stereomicroscope equipped with a Zeiss Axiocam 208 color camera (Carl Zeiss, Oberkochen, Germany). Wound areas were quantified using Fiji software (National Institutes of Health, Bethesda, MD, USA) and normalized to the initial wound size on day 0. To evaluate inflammatory monocyte/macrophage accumulation during healing, intravital multiphoton imaging was performed in CCR2-GFP mice (JAX 027619). Mice were anesthetized with isoflurane and imaged using a Leica SP8 DIVE multiphoton microscope (Leica Microsystems, Wetzlar, Germany). CCR2-positive inflammatory monocytes/macrophages were visualized by GFP fluorescence, whereas type I collagen fibers were visualized using second harmonic generation (SHG) imaging. The same wound region in each mouse was imaged repeatedly on day 0 and day 3 after wound induction. CCR2-positive-cell fluorescence intensity was quantified using Fiji software, and inflammatory-cell accumulation was expressed as the day 3/day 0 fluorescence-intensity ratio. Each experimental group included four mice (n = 4 per group).

## 3. Results

### 3.1 Schematic illustration of lyophilized EV preparation and in vivo wound-healing experimental design

A schematic overview of the experimental workflow used in this study is presented in Figure 1. MSCs were cultured under standard conditions, and EVs were isolated from MSC-conditioned medium. Isolated Evs were then subjected to lyophilization and stored at −80 °C to preserve their structural stability and biological activity. To evaluate the regenerative efficacy of lyophilized Evs in vivo, a circular ear wound model was established in CCR2-GFP mice. After wound generation, lyophilized Evs were topically administered to the injury. Wound-healing progression was monitored over time, and the EV-treated group was compared with the control group during the healing process.

**Figure 1.**
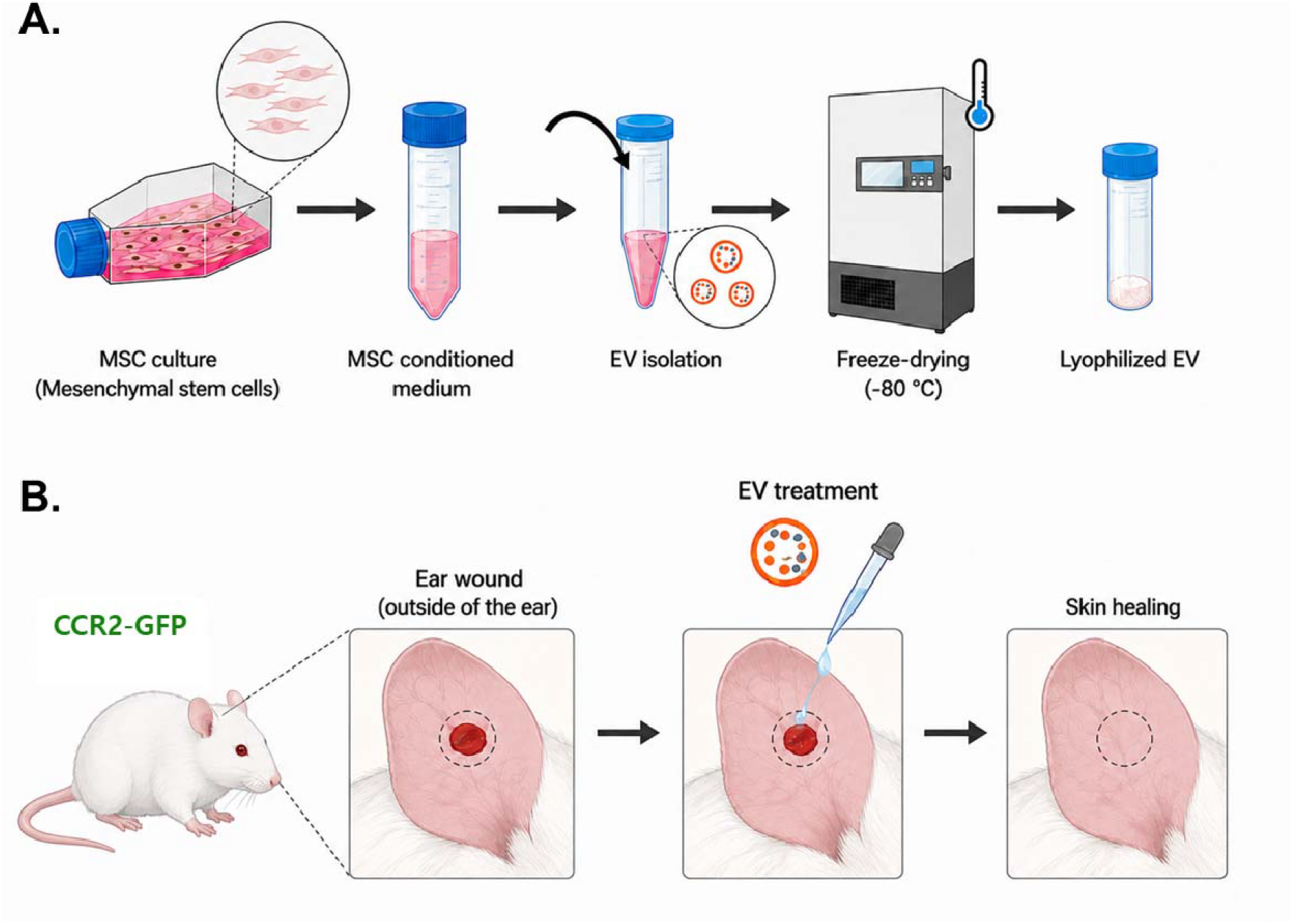
Schematic illustration of the experimental workflow for lyophilized EV preparation and in vivo wound-healing evaluation. Human epidural fat-derived mesenchymal stem cells (hEF-MSCs) were cultured under standard conditions, and extracellular vesicles (EVs) were isolated from conditioned medium. Isolated EVs were lyophilized and stored at −80 °C. For in vivo evaluation, a full-thickness ear wound was generated in CCR2-GFP reporter mice using a 1-mm biopsy punch. Lyophilized EVs were reconstituted and topically administered to the wound site. Wound-healing progression and inflammatory-cell responses were then assessed using macroscopic imaging and intravital multiphoton microscopy.

### 3.2 Characterization of hEF-MSCs

The morphological characteristics and stem cell properties of hEF-MSCs were evaluated before EV isolation. As shown in Figure 2A, cultured hEF-MSCs exhibited a typical spindle-shaped, fibroblast-like morphology under phase-contrast microscopy. The cells showed homogeneous growth patterns and maintained stable morphology during culture expansion, consistent with general MSC characteristics. To further confirm the MSC phenotype, surface marker expression was analyzed by flow cytometry. hEF-MSCs highly expressed the representative MSC markers CD90 (76.51%) and CD105 (74.87%), whereas expression of the hematopoietic marker CD45 was minimal (Figure 2B). These results confirmed that the cultured cells showed the characteristic immunophenotypic profile of MSCs and lacked substantial hematopoietic cell contamination. The multipotent differentiation capacity of hEF-MSCs was then assessed using trilineage differentiation assays. After lineage-specific induction, hEF-MSCs differentiated successfully into chondrogenic, adipogenic, and osteogenic lineages. Chondrogenic differentiation was confirmed by positive Alcian Blue staining, indicating proteoglycan-rich extracellular matrix formation. Adipogenic differentiation was verified by Oil Red O staining, which revealed intracellular lipid-droplet accumulation. Osteogenic differentiation was demonstrated by positive Alizarin Red S staining, indicating calcium deposition and mineralized matrix formation (Figure 2C). Collectively, these results confirmed that isolated hEF-MSCs retained the morphology, immunophenotype, and multipotent differentiation capacity characteristic of MSCs, supporting their suitability as a source for EV isolation and subsequent regenerative studies.

**Figure 2.**
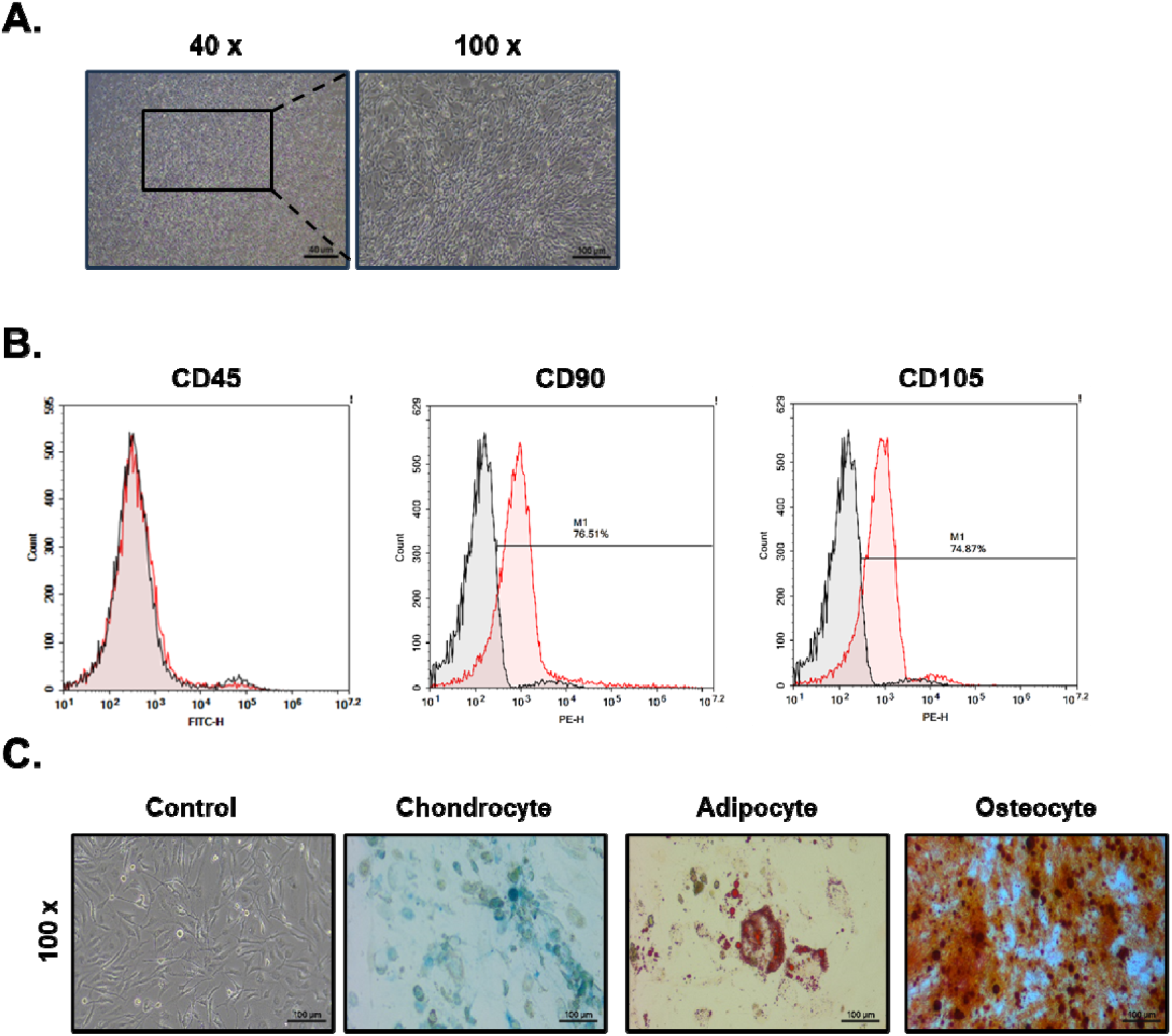
Characterization of human epidural fat-derived mesenchymal stem cells (hEF-MSCs). (A) Representative phase-contrast image showing the typical spindle-shaped morphology of cultured hEF-MSCs. Scale bar = 100 μm. (B) Flow cytometric analysis of MSC surface marker expression. hEF-MSCs showed positive expression of CD90 and CD105 and minimal expression of the hematopoietic marker CD45. (C) Trilineage differentiation potential of hEF-MSCs. Chondrogenic differentiation was confirmed by Alcian Blue staining, adipogenic differentiation by Oil Red O staining, and osteogenic differentiation by Alizarin Red S staining. Scale bars = 100 μm for chondrogenic and osteogenic differentiation and 40 μm for adipogenic differentiation.

### 3.3 Characterization of Lyophilized EVs

The structural and physicochemical characteristics of EVs before and after lyophilization were evaluated using TEM, NTA, and flow cytometry-based surface marker analysis. As shown in Figure 3A and B, both freshly isolated and lyophilized EVs exhibited a typical spherical, membrane-bound morphology under TEM. No apparent structural collapse or severe aggregation was observed after freeze-drying, indicating that lyophilization preserved the overall vesicular architecture. To further assess the effect of lyophilization on EV size distribution and particle concentration, NTA was performed. Fresh EVs had a mean particle size of 127.5 ± 1.9 nm and a concentration of 2.32 × 10^10^ particles/mL, whereas lyophilized EVs had a mean particle size of 126.6 ± 2.7 nm and a concentration of 2.34 × 10^11^ particles/mL (Figure 3C and D). These findings showed that lyophilization did not significantly alter EV size distribution or particle yield. In addition, EV surface marker expression was analyzed by bead-based flow cytometry. Both fresh and lyophilized EVs showed strong positive expression of the representative exosomal markers CD9, CD63, and CD81 (Figure 3E and F). These surface marker expression patterns were well preserved after freeze-drying and reconstitution, suggesting that lyophilized EVs retained their characteristic exosomal properties. Collectively, these results showed that lyophilization under the present experimental conditions effectively preserved EV morphology, particle-size distribution, concentration, and exosomal surface marker expression, supporting EV structural stability after storage at −80 °C.

**Figure 3.**
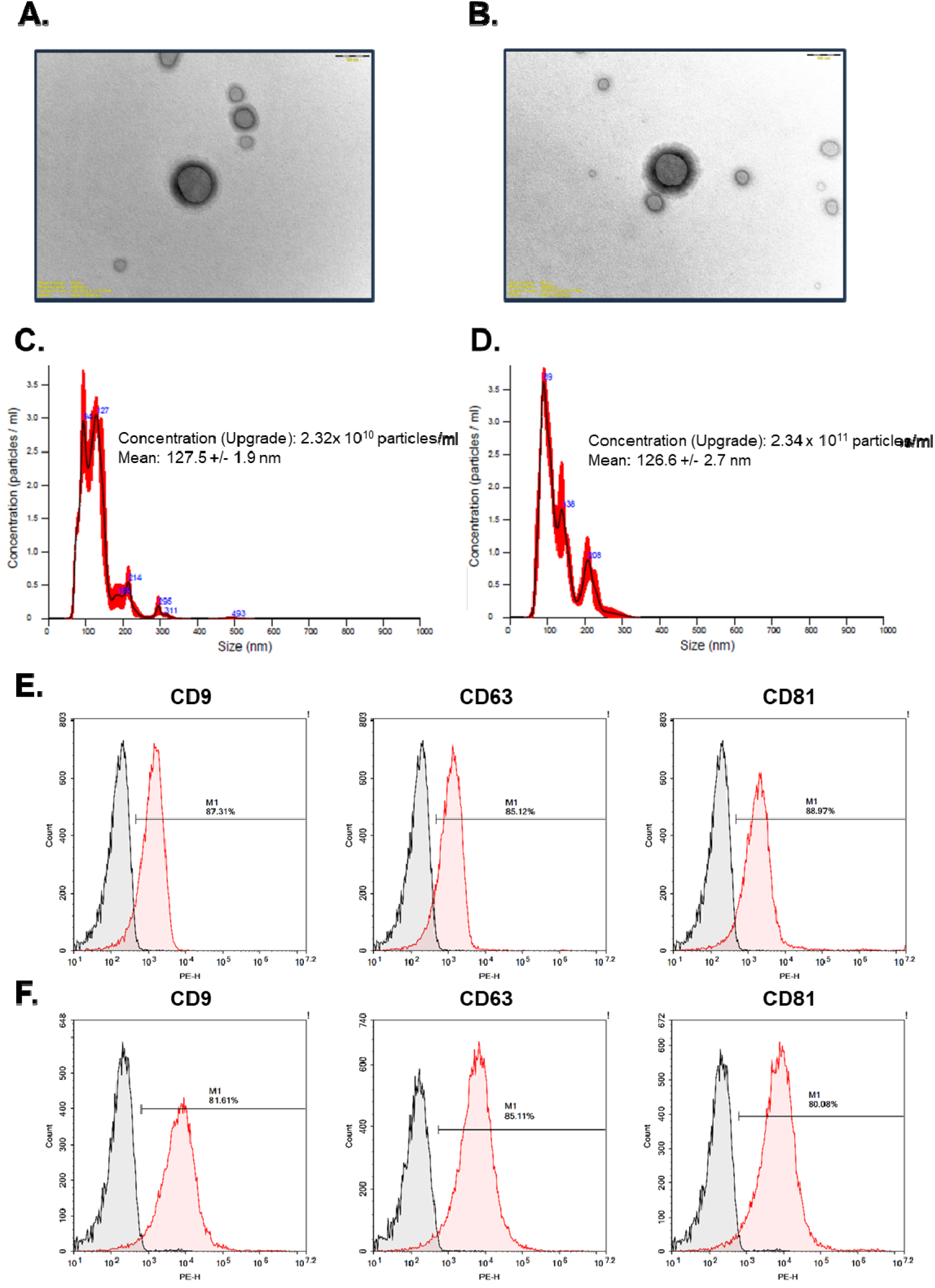
Characterization of extracellular vesicles before and after lyophilization. (A, B) Representative transmission electron microscopy (TEM) images of freshly isolated and lyophilized EVs after reconstitution. Both groups showed typical spherical vesicular morphology. Scale bars = 200 nm. (C, D) Nanoparticle tracking analysis (NTA) showing the particle-size distribution and concentration of fresh and lyophilized EVs. Fresh EVs exhibited a mean particle size of 127.5 ± 1.9 nm and a concentration of 2.32 × 10^10^ particles/mL, whereas lyophilized EVs exhibited a mean particle size of 126.6 ± 2.7 nm and a concentration of 2.34 × 10^11^ particles/mL. (E, F) Bead-based flow cytometric analysis of EV surface markers. Both fresh and lyophilized EVs expressed the representative exosomal markers CD9, CD63, and CD81, indicating preservation of EV characteristics after lyophilization.

### 3.4 Lyophilized EVs promote cutaneous wound repair through modulation of CCR2-positive inflammatory cells

To investigate the therapeutic potential of EVs in cutaneous wound healing, we employed an established mouse ear skin wound model combined with intravital imaging (15, 17). The ear skin model was selected because, unlike back skin, it undergoes minimal contraction during healing and therefore more faithfully recapitulates wound healing dynamics observed in humans (18). Full-thickness wounds were generated on the dorsal surface of the mouse ear using a 1-mm biopsy punch, followed by immediate topical application of lyophilized EVs resuspended in PBS. Control animals received an equivalent volume of PBS alone. Serial macroscopic imaging showed that EV-treated wounds underwent markedly accelerated closure relative to PBS-treated controls (Figure 4A and B). EVs have been widely implicated in the modulation of inflammatory responses (19-21).

**Figure 4.**
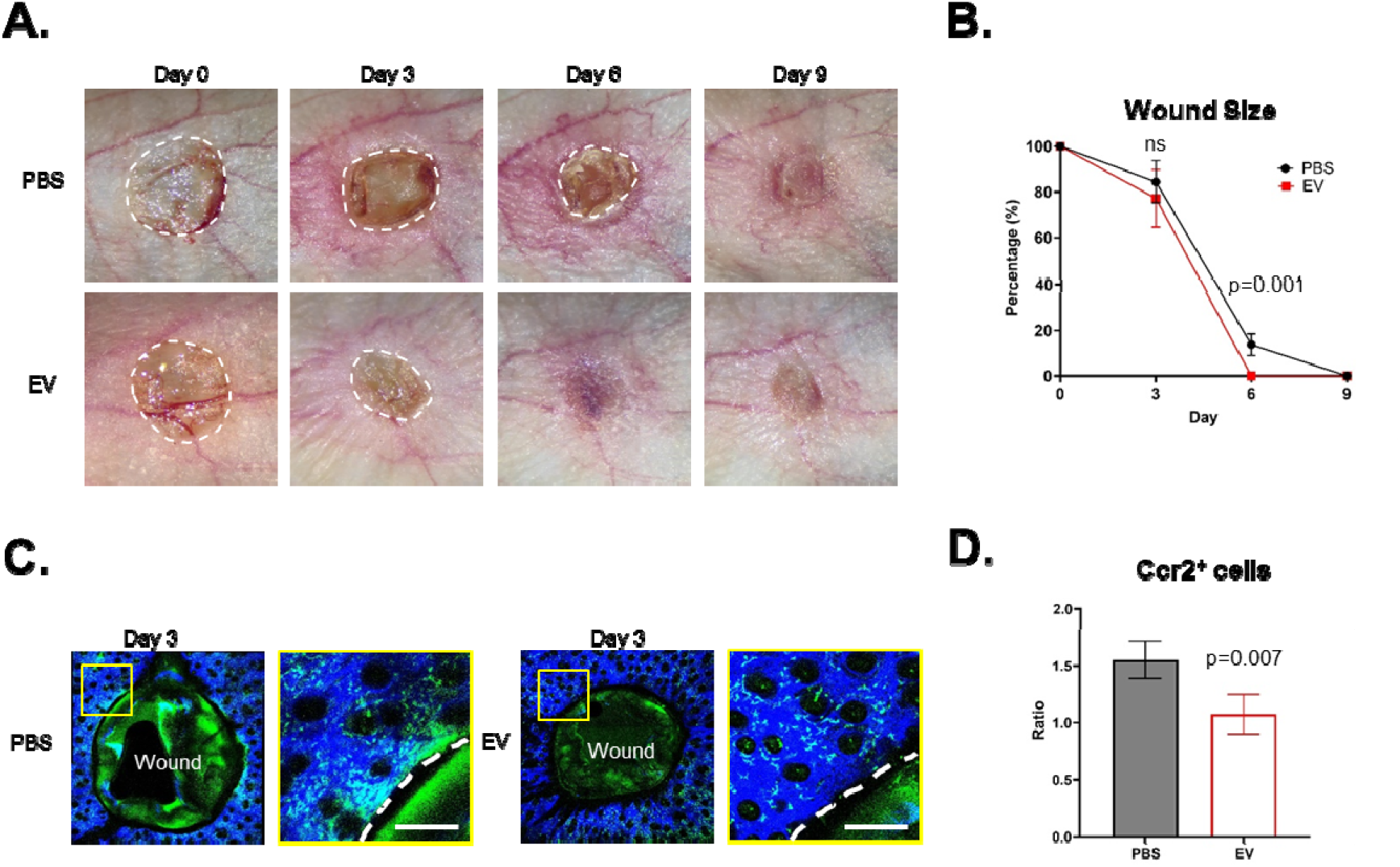
EVs accelerate wound healing by reducing inflammation in vivo. (A) Macroscopic images of ear skin wounds during healing after PBS or EV treatment. Full-thickness 1-mm punch biopsy wounds were generated on the dorsal ear skin of mice and imaged repeatedly at the indicated time points (days 0, 3, 6, and 9). White dashed lines indicate the wound boundary. n = 4 mice per group. (B) Quantification of wound size over time after PBS or EV treatment. Wound area was normalized to the initial wound size on day n = 4 mice per group. (C) Intravital images of inflammatory monocyte/macrophage accumulation at the wound site 3 days after wound induction. CCR2-positive inflammatory monocytes/macrophages are shown in green (CCR2-GFP), and type I collagen second harmonic generation (SHG) signal is shown in blue. Dashed lines indicate the wound boundary. n = 4 mice per group. Scale bars, 200 µm. (D) Quantification of CCR2-positive inflammatory monocyte/macrophage accumulation at the wound site in PBS-and EV-treated mice. The same mice were imaged repeatedly on day 0 and day 3 after wound induction, and the day 3/day 0 ratio was quantified. n = 4 mice per group.

Given that excessive or unresolved inflammation is a principal driver of impaired wound healing (22, 23), we next examined whether EV treatment altered inflammatory immune cell recruitment at the wound site. Previously, we established intravital imaging approaches to visualize CCR2-positive inflammatory monocytes in the skin of live CCR2-GFP reporter mice (24). Using this approach, we repeatedly imaged the same wound area immediately after injury and at day 3 post-injury and quantified the accumulation of CCR2-positive inflammatory immune cells during wound healing. Interestingly, EV-treated wounds showed a significant reduction in CCR2-positive cell accumulation compared with PBS-treated controls (Figure 4C and D). Collectively, these findings show that topical EV treatment accelerates wound closure while concurrently attenuating the accumulation of CCR2-positive inflammatory immune cells at the wound site, suggesting that EV-mediated immunomodulation contributes mechanistically to improved cutaneous tissue repair.

## 4. Discussion

EVs have emerged as promising cell-free therapeutic agents because of their ability to regulate tissue regeneration and modulate immune responses (2, 4). However, the clinical translation of EV-based therapeutics remains limited by challenges associated with long-term storage and the preservation of biological activity (5). In the present study, we showed that MSC-derived EVs retained their structural characteristics and therapeutic efficacy after lyophilization and storage at −80 °C. Furthermore, topical administration of lyophilized EVs significantly accelerated wound closure and reduced the accumulation of CCR2-positive inflammatory cells in a mouse ear wound model, suggesting a potential association between EV-mediated modulation of inflammatory cell recruitment and improved tissue repair.

Long-term preservation is a critical consideration for the clinical application of EV-based therapeutics (25). Conventional EV storage methods typically require continuous ultra-low-temperature storage and may result in gradual declines in vesicle stability and biological functionality during repeated freeze–thaw cycles. Lyophilization has emerged as a promising preservation strategy because it removes water while maintaining EV structure and integrity. In the present study, TEM, NTA, and exosomal surface marker analyses showed that lyophilized EVs retained their characteristic morphology, particle size distribution, concentration, and expression of CD9, CD63, and CD81 after storage at −80 °C. These findings are consistent with previous reports indicating that appropriately optimized lyophilization protocols can preserve EV integrity and facilitate their long-term storage and therapeutic application (26).

Beyond structural preservation, maintaining biological activity after lyophilization is essential for therapeutic applications. Although several studies have reported the physicochemical stability of freeze-dried EVs, evidence supporting the retention of their regenerative efficacy in vivo remains limited (27). Using a mouse ear wound model, we observed significantly accelerated wound closure in EV-treated animals compared with PBS-treated controls. Because the ear wound model shows minimal wound contraction and more closely resembles human wound healing than conventional dorsal skin wound models, these findings suggest that lyophilized EVs retain sufficient biological activity to promote tissue regeneration after storage. Our results support the feasibility of using lyophilized EV formulations for future translational and clinical applications.

Wound healing is a complex biological process involving coordinated inflammatory responses, angiogenesis, extracellular matrix remodeling, and tissue regeneration (28). Among these processes, regulation of inflammation is particularly important because excessive or prolonged inflammatory responses can delay healing and contribute to tissue damage. Previous studies have shown that MSC-derived EVs possess potent immunomodulatory properties through the delivery of bioactive proteins, lipids, and regulatory RNAs (29). Consistent with these observations, our intravital imaging analysis revealed that EV treatment significantly reduced the accumulation of CCR2-positive inflammatory cells at the wound site during the early stages of healing.

CCR2-positive monocytes/macrophages are key mediators of inflammatory responses after tissue injury and are rapidly recruited to damaged tissues through chemokine-dependent signaling pathways (30). Although transient recruitment of inflammatory monocytes is essential for host defense and tissue repair, excessive or prolonged accumulation can sustain inflammation and impair regenerative processes (31). In the present study, reduced accumulation of CCR2-positive cells in EV-treated wounds was associated with enhanced wound closure, suggesting that lyophilized EVs may facilitate tissue repair by modulating inflammatory cell recruitment. Although the precise molecular mechanisms underlying this effect remain to be elucidated, our findings support the hypothesis that EV-mediated immunomodulation contributes to the therapeutic benefits observed during wound healing.

Several limitations should be acknowledged in the present study. First, the molecular cargo responsible for the observed therapeutic effects was not investigated. Future studies examining EV-associated microRNAs, proteins, and signaling pathways may provide mechanistic insights into the regenerative properties of lyophilized EVs (32). Second, only a single storage condition (−80 °C) and wound model were evaluated. Additional studies comparing different storage durations and disease models would further strengthen the translational relevance of these findings (33). Finally, histological analyses of angiogenesis, collagen remodeling, and macrophage polarization were not performed and should be addressed in future investigations.

In conclusion, lyophilized MSC-derived EVs stored at −80 °C retained their structural integrity and biological activity. Topical administration of these EVs significantly accelerated wound healing and reduced CCR2-positive inflammatory-cell accumulation in vivo. These findings highlight lyophilized EVs as a practical and clinically applicable therapeutic platform for tissue regeneration and wound repair and emphasize immunomodulation as a key mechanism underlying their regenerative effects.

